# Enhancing microRNA activity through increased endosomal release mediated by nigericin

**DOI:** 10.1101/367672

**Authors:** Esteban A. Orellana, Loganathan Rangasamy, Srinivasarao Tenneti, Ahmed M. Abdelaal, Philip S. Low, Andrea L. Kasinski

## Abstract

The therapeutic promise of small RNA therapeutics (siRNAs, miRNAs) is not only limited by the lack of delivery vehicles, but also by the inability of the small RNAs to reach intracellular compartments where they can be biologically active. We previously reported successful delivery of functionally active miRNAs via receptor-mediated endocytosis^1^. This type of targeted therapy still faces one of the major challenges in the delivery field, endosomal sequestration. Here, a new method has been developed to promote endosomal escape of delivered miRNA. The strategy relies on the difference in solute contents between nascent endosomes and the cytoplasm: early endosomes are rich in sodium ions (Na^+^) while the intracellular fluid is rich is potassium ions (K^+^). Exploiting this difference through favoring the influx of K^+^ into the endosomes without the exchange for a osmotically active ion (Na^+^), results in an osmotic differential leading to endosome swelling and bursting. One molecule that is able to exchange K^+^ for an osmotically inactive hydrogen ion is the ionophore nigericin. Via generating an intramolecular miRNA delivery vehicle, containing a ligand, in this case folate, and nigericin we achieve escape of folate-RNA conjugates (e.g. FolamiRs) from their entrapping endosomes into the cytoplasm where they bind the RNA Induced Silencing complex (RISC) and activate the RNAi response.

## Introduction

The therapeutic prospect of miRNA replacement therapies relies on reintroducing small synthetic RNA molecules that mimic endogenous miRNAs in cells that have lost them with the final goal of restoring cellular pathways that could stop pathogenesis. However, a major constraint in the transition of miRNA mimics, and other small RNAs (small interfering RNA, siRNA; antisense oligonucleotides), into the clinic has been the lack of efficient delivery systems. Encapsulation of small RNAs, for instance, has had mixed results in the clinic with clinical trials showing toxicity and limited efficacy^2^. Some problems that come with the use of encapsulating delivery vehicles (e.g. liposomes, micelles, polymers) include delivery-associated toxicity, poor transfection efficiency, and nonspecific biodistribution^3^.

Our previous work shows that these vehicle-related defects may be overcome by eliminating the vehicle, attaching a miRNA to a folate ligand (FolamiR)^1^. This approach can deliver functionally active miRNAs to cancer cells that overexpress the folate receptor (FR) via receptor-mediated endocytosis with no signs of toxicity (maximum tolerated dose >26.64 mg/kg)^1^. Thus, in the absence of toxicity, the main factors limiting dosage would be two-fold: i) the internalization kinetics of the folate-ligand bound to its receptor (endocytosis) and ii) the rate by which the FolamiR can escape the endosome^4,5^.

We reasoned that since the internalization kinetics of FolamiRs are less amendable and already rapid^6–8^, the development and inclusion of an endosomal escape mechanism onto next-generation FolamiRs could help overcome the rate-limiting step of endosomal escape providing higher efficacy, and further reducing toxicity. Endosomal escape of small RNAs (and other biologics) has been recognized as a major hindrance in the path to translating RNA therapeutics into the clinic. For that reason efforts have been focused on developing strategies to enhance endosomal release and secure cytosolic delivery^9^. Proteins and peptide-based agents (i.e. cell penetrating peptides, CPPs) are often used to achieve cytosolic release of oligonucleotides^9^. For instance, the EB1 peptide, derived from penetratin, is a CPP with endosomolytic properties that facilitates transport of active siRNAs into cells^10^. Similarly, the inclusion of influenza-derived fusogenic peptide diINF-7 has been shown to enhance endosomal escape of siRNAs when complexed with lipid carriers^11^. However, the use of peptides is often toxic and may contribute to increased immunogenicity^12^, which is already a major concern in RNA therapeutics. Another approach to improve endosomal release of nucleic acids is the use of “proton sponges” that can promote endosomal swelling by inducing counterion uptake that leads to increased osmotic pressure^13^. The practicality of using this strategy in the clinic is dampened due to the large concentration of compound needed to achieve efficient endosomal bursting^13^. Photochemical internalization (PCI) is another alternative to increase endosomal escape of siRNAs, but this approach is greatly limited to specific sites where light can be used to trigger endosome lysis, which reduces its practical use in the clinic^14^.

We recently described a novel method to facilitate the escape of biologics from entrapping endosomes and in that study provided preliminary evidence for the potential use of this method to improve the escape of small RNAs from endosomes^15^. The strategy relies on the difference in solute contents between nascent endosomes and the cytoplasm: endosomes are rich in sodium ions (Na^+^) while the intracellular fluid is rich is potassium ions (K^+^). The method exploits this difference by using a small molecule, nigericin^16,17^, that exchanges K^+^ for osmotically inactive proton (H^+^) to achieve balance in charge without release of a compensatory solute (Na^+^) thus causing a osmotic differential that leads to endosome swelling and bursting^15^.

Herein, we show that this approach is useful to overcome endosomal entrapment of small RNAs delivered by folate. The data presented here suggest that ligand-targeted delivery of nigericin into endosomes can facilitate the escape of RNA cargo (e.g. miRNAs, siRNAs) from their entrapping endosomes, helping the small RNAs to become available in the cytoplasm, engage the RNA Induced Silencing complex (RISC) and improve their RNAi activity (see proposed model in Figure 1). Developing a highly specific delivery platform (i.e. FolamiRs) with increased endosomolytical properties could be useful towards achieving significant therapeutic effects at low concentrations and without unwanted toxicity for future *in vivo* applications.

**Figure 1:**
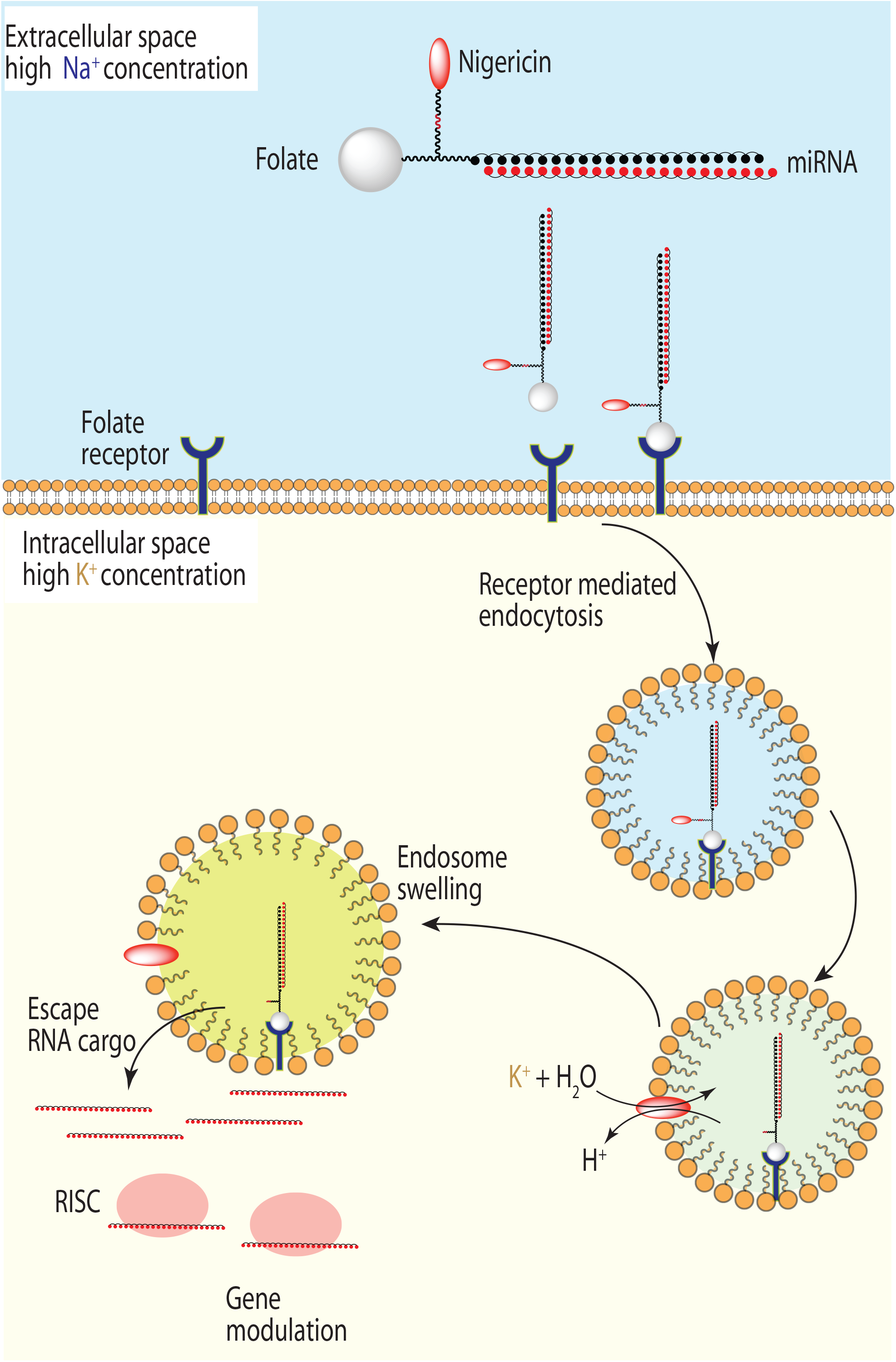
Proposed mechanism of action of endosomal escape of RNA cargo mediated by ligand-targeted delivery of nigericin. Internalization of folate-nigericin-RNA (Fol-Nig-RNA) conjugates is mediated by endocytosis. The nascent vesicle is rich in extracellular fluid with a high concentration of Na^+^ (blue shading), while the intracellular space contains high levels of K^+^ (yellow shading). Upon ligand-targeted delivery, nigericin, a K^+^/H^+^ antiporter, translocates to the endosomal membrane (following reduction of the di-sulfide bond, red linker) causing an influx of K^+^ (green shading) in exchange for H^+^. The exchange of K^+^ for the osmotically inactive H^+^ leads to build up of osmotic pressure which causes endosomal swelling and release of RNA cargo into the cytosol.

## Results

### Folate-nigericin-miRNA conjugation and stability in serum

Since nigericin needs to dissociate from the folate carrier and localize to the endosomal membrane to induce endosome swelling (exchange intracellular K^+^ for H^+^), an intramolecular disulfide linkage was incorporated between nigericin and the folate-RNA conjugate (Figure 2A). Folate-nigericin-miRNA conjugation was achieved as previously described^1^ by using click chemistry^26^ and conjugation was verified by polyacrylamide gel electrophoresis (PAGE) analysis (Figure 2B) and matrix-assisted laser desorption/ionization (MALDI) spectral analysis (not shown). Disulfide linkage reduction was verified by PAGE analysis by a band size shift upon tris(2- carboxyethyl)phosphine (TCEP) treatment (Figure 2B). To test the stability of the disulfide linkage, Fol-Nig-34a compounds were exposed to 50% serum. No signs of Fol-Nig-34a reduction were observed for more than four hours (Figure 2C) leaving open the possibility that premature reduction upon systemic delivery could occur after 4 h. Our previous results using FolamiRs in vivo show that less than four hours is enough time for the folate conjugates to localize and be internalized into the intended tissues^1^.

**Figure 2:**
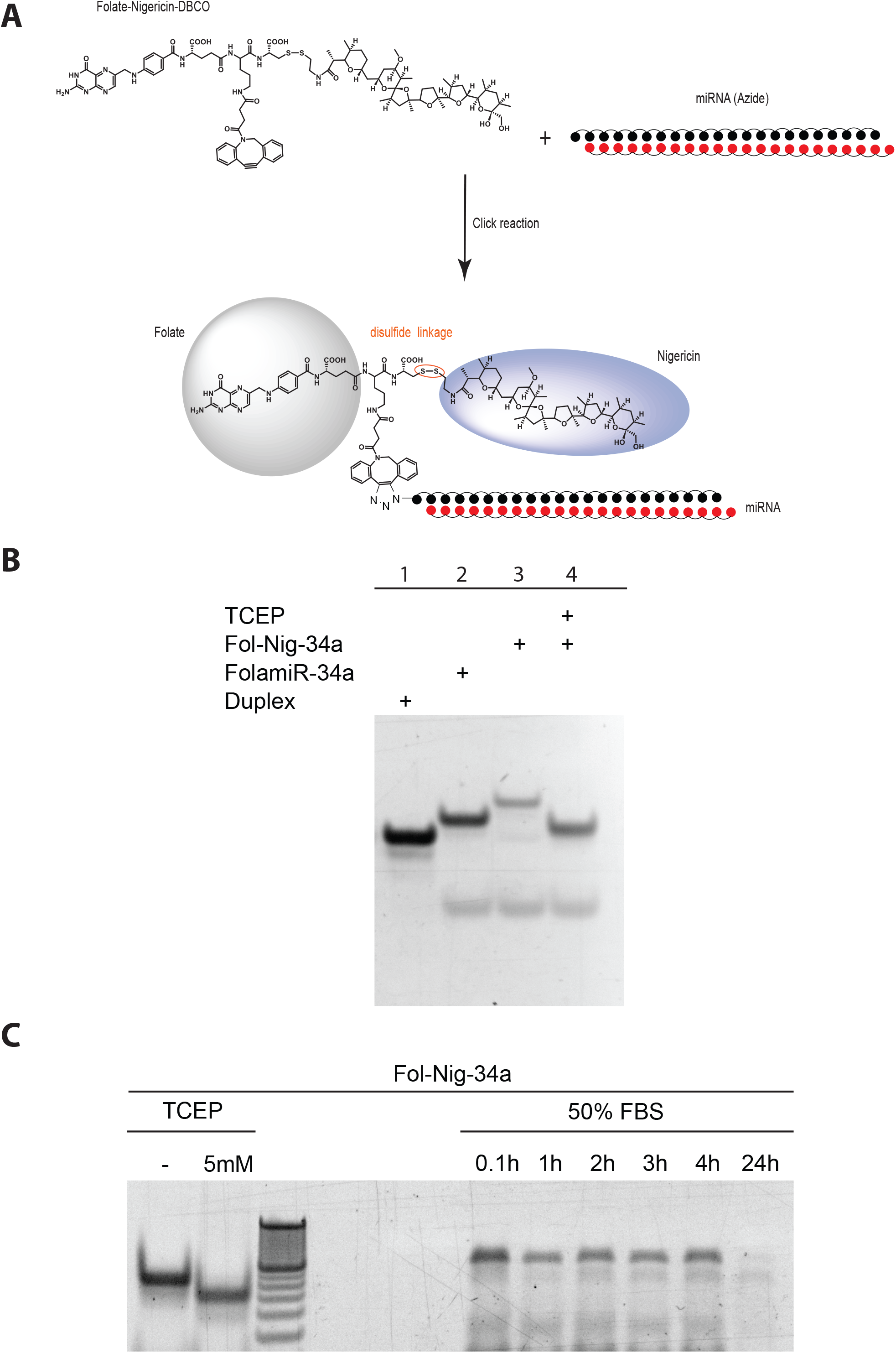
Evaluation of folate-nigericin-miRNA conjugation and stability in serum. **A)** Scheme of folate-nigericin-miR34a (Fol-Nig-34a) conjugation. **B)** Gel red stained polyacrylamide gel. Each lane was loaded with 200 pmol of unconjugated oligo (Duplex) or folate-miRNA. “+” indicates the presence of a particular ligand conjugated to miR-34a duplex. PAGE results suggest successful conjugation of folate-miRNA compounds as visualized by the shift in mobility. The band shift in lane 4 indicates reduction of the disulfide bond between nigericin and folate-miRNA. TCEP, reducing agent tris(2-carboxyethyl)phosphine. **C)** Gel red stained polyacrylamide gel following resolution of folate-nigericin-miRNA after 50% serum exposure for different periods of time. TCEP (5mM) is included as a positive control.

### Folate-nigericin-miRNA conjugate uptake is mediated by the folate receptor

Due to the high affinity of the ionophore nigericin for cellular membranes it was important to verify that uptake of Fol-Nig-miRNA conjugates was mediated by the FR. For that purpose, expression of the FR in MB-231 cells and KB cells was verified by flow cytometric analyses as previously reported^1,27^ using a folate-cyanine 5 conjugate (Fol-Cy5; Figure 3A). Next, to visualize cellular uptake of miR-34a in the absence or presence of nigericin, an Atto-647N fluorophore was attached to the 5′ end of miR-34a-5p strand generating Fol-34a-Atto647N and Fol-Nig-34a-Atto647N. Specificity of cellular uptake of Fol-34a-Atto647N or Fol-Nig-34a-Atto647N conjugates was conducted in the presence or absence of folate glucosamine as a competitor (Figure 3B,C). Both conjugates were taken up by MB-231 and KB cells only in the absence of folate glucosamine suggesting that the uptake is FR mediated. Furthermore, minimal uptake in the absence of folate conjugation (gymnotic uptake) was observed (Figure 3D, duplex miR-34a-Atto647N). Taken together these results suggest that Fol-Nig-miRNA uptake is mediated by the FR.

**Figure 3:**
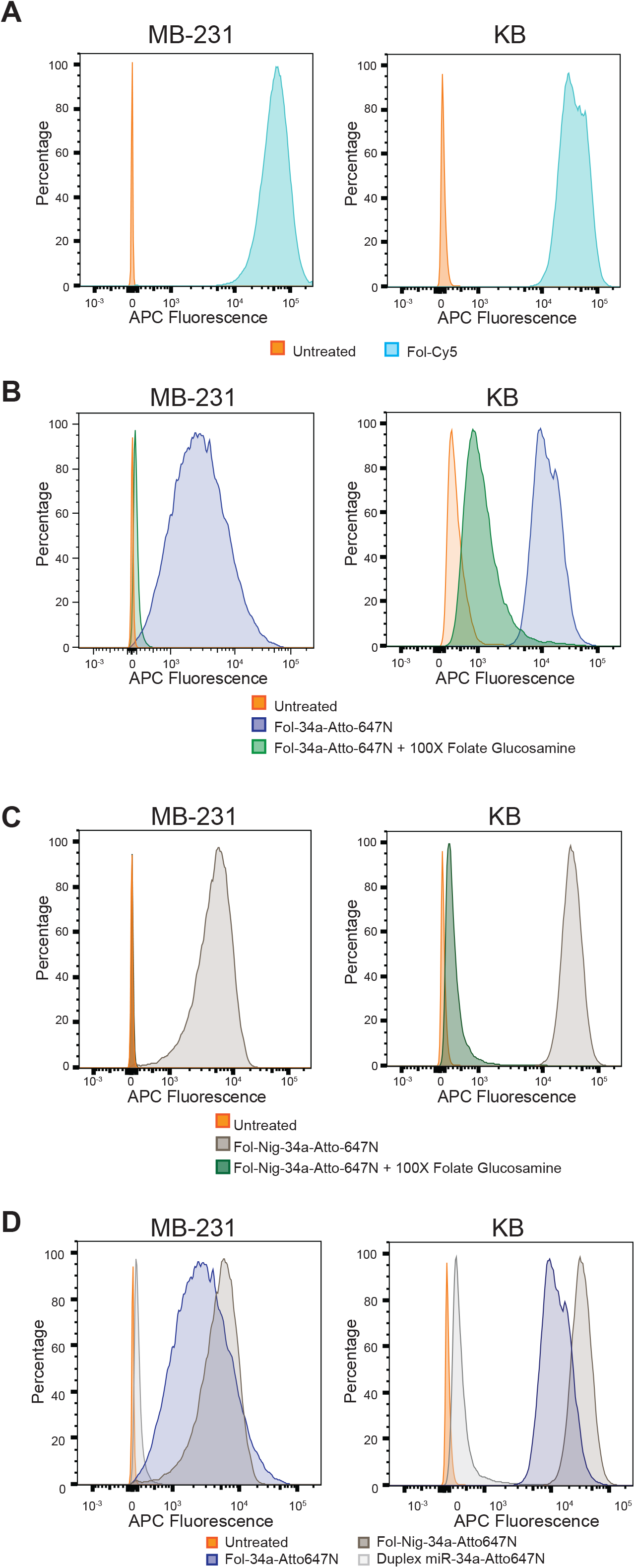
Folate-nigericin-miRNA conjugate uptake is mediated by the folate receptor. **A)** Confirmation of folate receptor expression in MB-231 breast cancer cells and in KB carcinoma cells. Histograms represent overlaid flow cytometry data as a percentage of unstained and folate-cyanate 5 (Fol-Cy5) cells. Displacement of Fol-34a-Atto647N **(B)** or Fol-Nig-34a-Atto647N **(C)** binding from MB-231 and KB cells (50 nM, 4ºC) in the presence of 100-fold excess of folate-glucosamine conjugate. Histograms represent overlaid flow cytometry data as a percentage of unstained, and Fol-34a-Atto647N or Fol-Nig-34a-Atto647N stained cells. **(D)** Folate-nigericin-miRNA conjugate uptake is not mediated by gymnosis. MB-231 or KB cells were treated with the Fol-34a-Atto647N, Fol-Nig-34a-Atto647N, or with duplex miR-34a-Atto647N in the absence of transfection reagent. Histograms represent overlaid flow cytometry data as a percentage of unstained, Fol-34a-Atto647N, Fol-Nig-34a-Atto647N or miR-34a-Atto647N stained MB-231 and KB cells (50 nM, 4ºC).

### Ligand-targeted delivery of nigericin causes endosomal release of miR-34a

To track the release of cargo from endosomes, cyanide 5 (Cy5) was delivered either with a folate-only or a folate-nigericin carrier, and the fluorescent cellular distribution was used as a surrogate for cargo localization. For that purpose, FR positive MB-231 cells were incubated with 50 nM of Fol-Cy5 or Fol-Nig-Cy5 and imaged every two min for 1 h. Representative images of the cells at 15 min following treatment are presented in Figure 4. The intracellular distribution of Cy5 was noticeably different in the absence of nigericin. Cells treated with Fol-Cy5 show fluorescent signal in their cell membrane and individual punctate spots through the cytoplasm (Figure 4A) indicative of endosomal localization^8^. In contrast, cells treated with Fol-Nig-Cy5 show perinuclear distribution as string-like dispersions with no clearly distinguishable punctate spots (Figure 4A).

**Figure 4:**
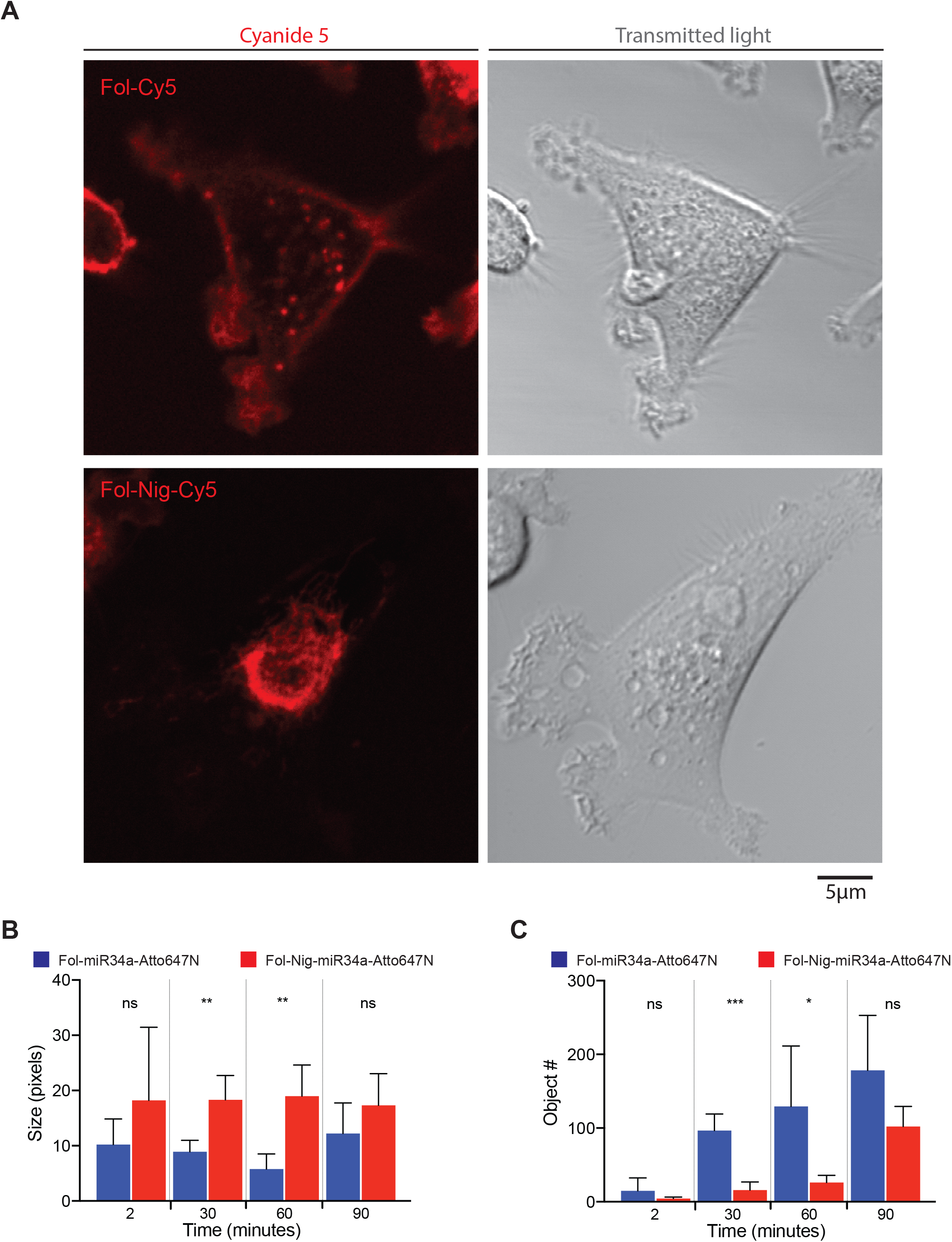
Ligand-targeted delivery of nigericin promotes cargo escape form endosomes. **A)** Representative confocal images of MB-231 cells treated with Fol-Cy5 or Fol-Nig-Cy5 (50 nM) at 15 min post treatment. Comparison of fluorescent object size **B)** and object number **C)** between Fol-34a-Atto647N or Fol-Nig-34a-Atto647N treated KB cells at different time points. (n = 4, error bars: mean ± s.d.). Segmentation and quantification of subcellular vesicles was performed using the segmentation and quantification of subcellular shapes (Squassh) protocol^19,20^. Statistical analysis performed with Student’s T-test,*, P < 0.05; **, P < 0.01; ***, P < 0.001; ns, not significant.

Next, to monitor the localization of miR-34a in the absence or presence of nigericin, FR positive KB cells were incubated with 50 nM of Fol-34a-Atto647N or Fol-Nig-34a-Atto647N and imaged every two min for 3 h. As early as 30 min after treatment noticeably larger vesicles were observed in the cells treated with Fol-Nig-34a-Atto647N conjugate compared to the vesicles of cells treated Fol-34a-Atto647N (Figure 4B). Interestingly the number of fluorescent punctate detected in cells treated with Fol-Nig-34a-Atto647N is reduced compared to cells treated with Fol-34a-Atto647N (Figure 4C), which could suggest that miR-34a-Atto647N delivered with the folate-nigericin carrier is not entrapped inside of endosomes. Taken together, these data suggest that ligand-targeted delivery of nigericin promotes escape of cargo from entrapping endosomes.

### Ligand-targeted delivery of nigericin promotes miR-34a targeting and engagement in RISC

To validate that nigericin can mediate endosomal escape, we used a functional assay to monitor RNAi activity using miR-34a. Luciferase reporter assays were performed in MB-231 cells that express a reporter for miR-34a^1^. Renilla level in these cells is inversely correlated with miR-34a activity. These cell-based experiments indicated a rapid reduction in luciferase activity in Fol-Nig-34a treated cells as soon as 18h post treatment (reaching ∼40% knockdown) and after 48h reaching 85% (Figure 5A). This reduction in luciferase activity did not occur via gymnosis (see Figure 5A, gray bars) nor was it observed in Fol-Nig-NC treated cells suggesting that the effect is specific to the presence of miR-34a and not due to a side effect caused by potential nigericin mediated toxicity. To eliminate that possibility, we conducted cellular toxicity assays and the results show that the first signs of toxicity related to nigericin compounds (folate-nigericin) can be visualized at concentrations >1000nM (Figure 5B), well above the 50 nM concentration used for the functional assays. Next, we reasoned that since RNA-induced silencing complex (RISC) programming is triggered by the appearance of double stranded RNA in the cytoplasm (i.e. miRNA) and Ago proteins localize diffusely in the cytoplasm and nucleus^28,29^ loading of miR-34a-5p into Ago would be indicative of cytosolic release. We first tested if Fol-Nig-34a activity is mediated by Ago proteins by measuring cell proliferation when Ago2 is knocked down (Figure 5C). The results show that Fol-Nig-34a only reduces cellular proliferation when Ago2 is present, indicating that Fol-Nig-34a cellular effects are mediated in part by the RNAi pathway. These results were also reproducible using a Fol-Nig-siRNA conjugate against Ras homolog gene family, member A (siRhoA), a known target that affects cell proliferation in MB-231 breast cancer cells^18^ (Figure 5D). To further test the hypothesis that nigericin facilitates the release of RNA cargo into the cytoplasm, we performed RNA immunoprecipitation (RIP) assays by immunoprecitating Argonaute (Ago) proteins^21,22^ in cells treated with FolamiR-34a or Fol-Nig-34a (Figure 5E). The results show that miRNA-34a-5p is more abundant in Ago RIP following folate-nigericin delivery (∼6 fold) compared to delivery using the folate-only conjugation (Figure 5F, compare blue bar to red bar in miR-34a qRT lanes). Ago did not pull down RNU6B, as expected, but was associated with the positive control *let- 7b* validating the assay. Taken together, these results support the hypothesis that partial decrease in luciferase activity is RNAi mediated and that the inclusion of nigericin in the folate carrier can help miR-34a escape from endosomes and become available in the cytoplasm to associate with Ago.

**Figure 5:**
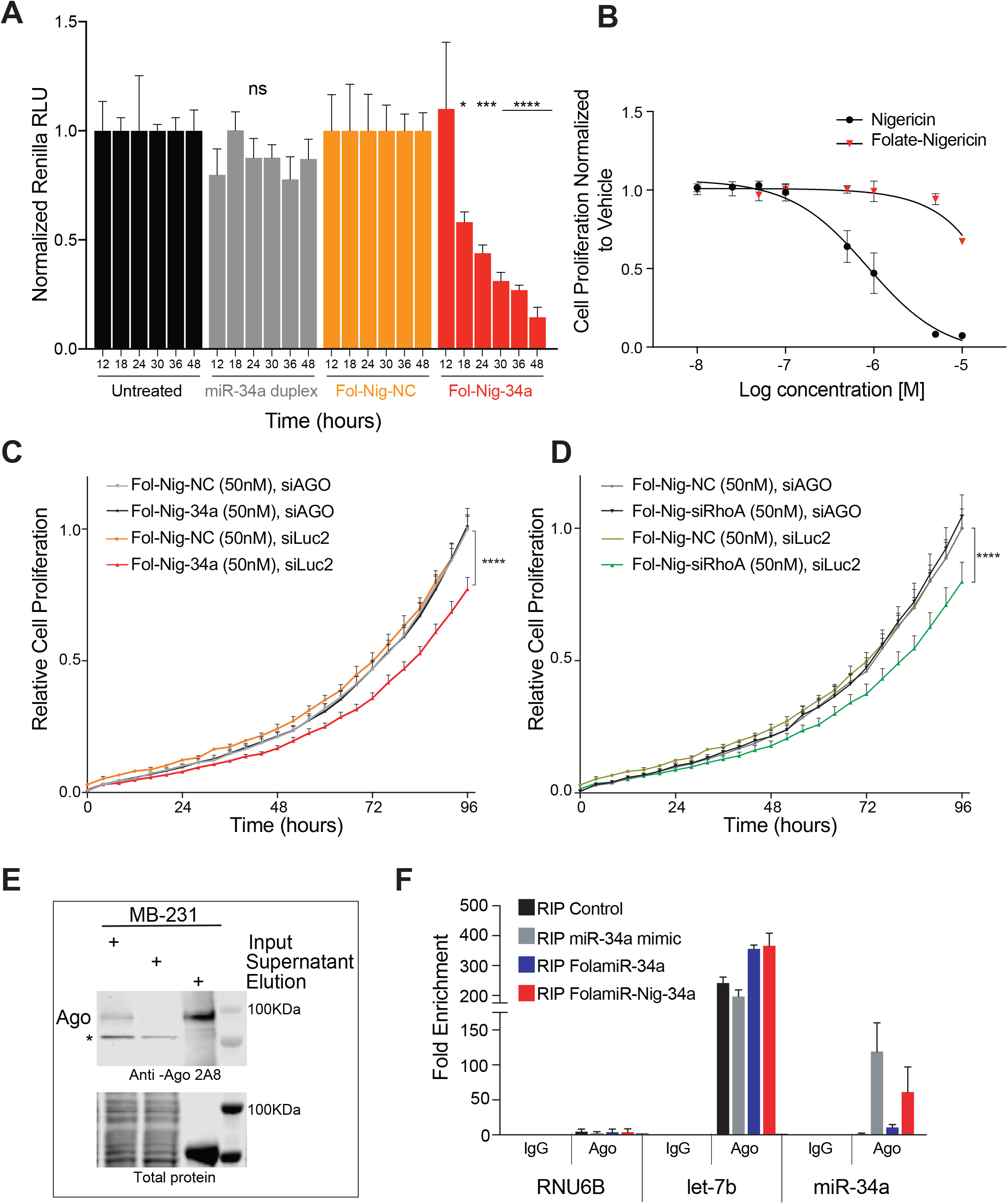
Nigericin mediates endosomal escape of miRNA cargo. **A)** Effect of Fol-Nig-34a on miR-34a sensor silencing in cells in culture. Fol-Nig-34a data points were normalized to Fol-Nig-NC (negative control: scrambled miRNA). Untransfected miR-34a duplex treated cells were normalized to untreated cells. Each experiment corresponds to n=3, 4 technical replicates per treatment, 50 nM, statistical analysis performed with one-way ANOVA and Bonferroni correction, (*, P < 0.05; ***, P < 0.001; ****, P < 0.0001). **B)** Effect of free folate and folate-nigericin on MB-231 cell growth. Data points were normalized to vehicle (DMSO). Error bars: mean ± s.d. n= 6. **C & D)** Cell proliferation of MB 231 cells transfected with the indicated siRNA (siAGO or control, siLuc) and treated with the indicated folate-nigericin conjugate, **C)** Fol-Nig-34a, and **D)** Fol-Nig-siRhoA. Experiments were conducted in replicates of 6, statistical analysis performed with two-way ANOVA and Bonferroni correction (****, P < 0.0001). **E)** Verification of Ago pulldown for Ago RIP experiments. * nonspecific band. **F)** RT-qPCR results following Ago RIP from cells treated with FolamiR-34a or Fol-Nig-34a (100 nM). Transfected miR-34a mimic was used as a delivery control (6 nM). RNU6B: not targeted control, *let-7b*: positive control. Experiments were repeated twice. Error bars: mean ± s.d.

### Inclusion of nigericin in the folate carrier enhances miRNA activity

To test the hypothesis that the incorporation of nigericin in the folate carrier can lead to enhanced miR-34a activity in culture time-response studies were conducted comparing FolamiR-34a and Fol-Nig-34a. The results show a difference in luciferase repression between FolamiR-34a (∼30% reduction in Renilla activity) and Fol-Nig-34a (∼85% reduction in Renilla activity, Figure 6A) that could indicate that FolamiR-34a gets entrapped inside endosomes by folate-targeted endocytosis. This could also explain why miR-34a delivered by folate alone is not abundant in Ago RIP (Figure 5F). Similarly, delivery of siRhoA by folate-nigericin results in higher anti-proliferative activity compare to delivery using the folate-only carrier (Figure 6B).

To verify that the released miR-34a can repress endogenous targets, MB-231 cells were exposed to FolamR-34a or Fol-Nig-34a and surface expression of PD-L1, a miR-34a target was monitored by flow cytometry. Only in the presence of Fol-Nig-34a was PD-L1 downregulated (Figure 6C). The effect observed was nearly identical to downregulation of PD-L1 following transfection of miR-34a mimics. Importantly, there was no reduction in PD-L1 in FR negative A549 cells (Figure 6D), unless cells were transfected with miR-34a mimics, suggesting dependency of the FR on this response. This dependency was verified in MB-231 cells, where excess folate-glucosamine mitigated the effect (Figure 6E). Taken together, these results support the hypothesis that rapid release of miRNA cargo from endosomal vesicles mediated by nigericin can improve miR-34a activity in culture.

**Figure 6:**
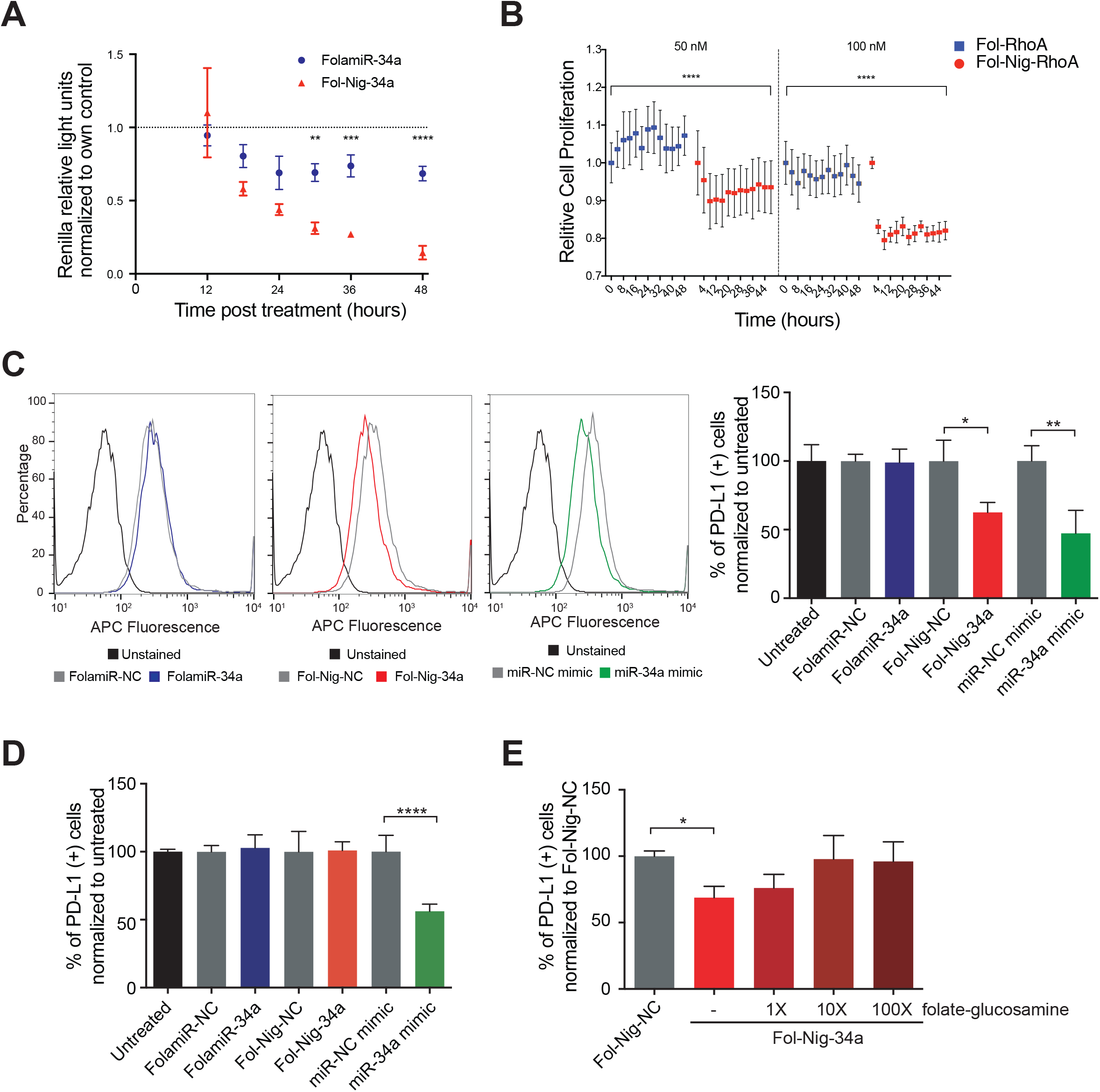
Incorporation of nigericin into the folate carrier enhances miRNA activity *in vitro*. **A)** Targeted silencing of the miR-34a Renilla sensor using Fol-Nig-34a in cells in culture. Error bars: mean ± s.d. Each experiment corresponds to n=3, 4 technical replicates per treatment. **B)** Effect of cellular proliferation in the presence of siRhoA conjugated to folate-only or folate-nigericin carrier. Statistical analysis performed with two-way ANOVA and Bonferroni correction, (**, P < 0.01; ***, P < 0.001; ****, P < 0.0001). **C)** Flow cytometry analysis of FR(+) MB-231 cells treated with the indicated folate-conjugate (100 nM), or transfected with the same dose of the indicated miRNA mimics. Representative assay on left. Quantified data from three biological replicates on right. **D)** Quantified flow cytometry data from three biological replicates of A549 cells treated with the indicated folate-conjugate (100 nM), or transfected with the same dose of the indicated miRNA mimics. Statistical analysis performed with one-way ANOVA and Bonferroni correction, (*, P < 0.1; **, P < 0.01; ***, P < 0.001; ****, P < 0.0001).

## Discussion

Development of an effective delivery strategy for targeted, tissue-specific delivery of small RNAs can be achieved with the use of small ligands. The addition of a ligand to a small RNA promotes its cellular uptake and trafficking, and although is a key advantage to achieve the required specific biodistribution, it often suffers from a major limitation, endosomal entrapment. Upon binding to the membrane receptor, these ligands and the associated RNA cargo are internalized by receptor-mediated endocytosis and loaded into early endosomes. The contents of the endocytic vesicles are then sorted and trafficked intracellularly en route to the lysosome, which is where the nucleic acid cargo will be degraded or are recycled back to the extracellular space. To avoid these therapeutically limiting effects, the ligand-RNA conjugates need to escape from the endosome into the cytoplasm where they can engage RISC and modulate gene expression. Thus, the clinical application of small RNA therapeutics requires the development of delivery strategies that not only can direct tissue-specific internalization, but also effectively traffic the small RNA to the cytoplasm reaching biologically relevant doses.

Our previous research showed that it is possible to deliver a miRNA mimic to cancerous cells by simply conjugating the passenger strand to a folate ligand with no signs of toxicity at the therapeutic dose tested^1^. Because folate conjugates are internalized via endocytosis, endosomal sequestration could be a limiting step in achieving significant clinical effects. We reasoned that the inclusion of an inter- molecular endosomal escape mechanism could ensure robust cytosolic delivery of the RNA cargo and lead to enhanced miRNA activity. This approach relies on the use of a small molecule, nigericin, that upon cellular internalization gets cleaved from the folate carrier and localizes to endosomal membrane where it exchanges K^+^ and water for osmotically inactive proton (H^+^) causing endosomal swelling^15^. To be effective, nigericin release has to occur in a controlled manner and only within the endosomes. Our preliminary evidence suggest that folate-nigericin-miRNA molecules are stable in 50% serum for at least four hours, which according to our previous observations would be enough time for the conjugate to be internalized into the target cells^1^. Premature cleavage and release of nigericin in circulation is a concern and further in vivo experiments will be needed to evaluate this possibility.

The evidence presented here supports that the new method for endosomal release of small RNAs can be advantageous therapeutically. The aims of this strategy are to decrease the dosage of mimics needed to achieve effective gene modulation and minimize unwanted toxicity. The data support that ligand-targeted delivery of nigericin into endosomes can facilitate the escape of RNA cargo (e.g. miRNAs, siR- NAs) from their entrapping endosomes, helping the small RNAs to become available in the cytoplasm, engage the RNA-Induced Silencing complex (RISC) and improve their RNAi activity. Thus, it is possible that with the help of nigericin, therapeutically relevant small RNAs could be delivered at low quantities and still show effective targeting. The most clinically advanced ligand-siRNAs, GalNAc–siRNA conjugates^30–34^, have shown mixed results and often require very high doses to achieve a significant biological effect. It is too early to know if the high doses of these oligonucleotides will have deleterious effects in the patients. Perhaps, achieving lower effective doses of therapeutic RNAs could be the key to avoid in vivo toxicity.

We anticipate that since nigericin is a small molecule, nontoxic, and easy to conjugate to a small RNA using a variety of self-cleavable linkers it will be easy to translate into the clinic. The challenge of the field of small RNA delivery is to continue to design and optimize delivery mechanism at all stages, from the delivery process, to cytosolic release, to uptake into RISC and mRNA cleavage such that small RNAs can be delivered to the intended tissues at low effective doses in the absence of unwanted toxicity. There is scientific awareness about the potential of miRNA therapeutics, but with the lack of effective cytosolic release strategies small RNA therapeutics are predetermined to bottleneck in the clinic. Although these molecules have yet to reach in vivo experiments, the rigorous understanding of the safety and efficacy profiles of this new delivery and release platform could eventually enable the transition of ligand-RNA conjugates from the bench to the bedside.

## Materials and methods

### Cell culture

MDA-MB-231 cells (MB-231, HTB-26), KB carcinoma cells (CCL-17), and A549 lung epithelial carcinoma cells (CLL-185), mycoplasma-free as determined by testing for mycoplasma contamination via MycoAlert Mycoplasma Detection Kit (Lonza), were grown in RPMI 1640 medium, no folic acid (Life Technologies) supplemented with 10% fetal bovine serum (Sigma), penicillin (100 U/mL), and streptomycin (100 ug/mL) (HyClone, GE Healthcare Life Sciences), and maintained at 37°C in 5% CO_2_. Authentication of MB-23, KB, and A549 cells was performed using short tandem repeat profiling [American Type Culture Collection (ATCC)].

### Preparation of pyridyldisulfide amide derivative of nigericin

Folate-ethylediamine (EDA), folate-cys and folate-dybenzocyclooctyne (DBCO) synthesis was performed as previously described^1^. Folate-nigericin synthesis was performed as described in^15^, nigericin free acid (0.035 mmol), Py-SS-(CH_2_)_2_NH_2_ (0.052 mmol), hexafluorophosphate azabenzotriazole tetramethyl uronium (HATU; 0.052 mmol), and N,N-Diisopropylethylamine (DIPEA; 0.069 mmol) were dissolved in anhydrous CH_2_Cl_2_ (2.0 mL) and stirred under argon at room temperature overnight. Progress of the reaction was monitored by liquid chromatography– mass spectrometry (LC-MS). After complete conversion of nigericin free acid, the crude reaction mixture was subjected to purification by R-HPLC, (mobile phase A = 10 mM ammonium acetate, pH = 7; organic phase B = acetonitrile; method: 0% B to 100% B in 35 min at 13 ml/min) and furnished nigericin-SS-amide derivative 55% yield. LC-MS (A = 10 mM ammonium bicarbonate, pH = 7; organic phase B = acetonitrile; method: 0% B to 100% B in 15 min) RT=9.15min(M+NH_4_^+^ =910.5)

### Preparation of folate-nigericin conjugate

Folate-nigericin synthesis was performed as previously described^15^: DIPEA was added dropwise to a stirred solution of folate-Cys (0.004 g, 0.007 mmol, 1.5 eq.) and pyridyldisulfide amide derivative of nigericin (0.004 g, 0.005 mmol, 1.0 eq.) in DMSO. The reaction mixture continued to be stirred at room temperature. Progress of the reaction was monitored by LC-MS. After complete conversion of folate-nigericin, the crude reaction mixture was purified by RPHPLC, (mobile phase A = 10 mM ammonium acetate, pH = 7; organic phase B = acetonitrile; method: 0% B to 40% B in 35 min at 13 ml/min) and furnished folate-nigericin 65% yield. LC-MS (A = 10 mM ammonium bicarbonate, pH = 7; organic phase B = acetonitrile; method: 0% B to 100% B in 7 min) RT = 4.0 min (M +H^+^ = 1440.0).

### Preparation of folate-nigericin-DBCO

Folate-nigericin-DBCO synthesis was performed as previously described^15^: N,NDiisopropylethylamine (DIPEA; 0.0013 g, 0.010 mmol, 1.5 eq.) was added dropwise to a stirred solution of folate-nigericin (0.010 g, 0.0069 mmol, 1 eq.) and N-Hydroxysuccinimide (NHS)-DBCO (0.003 g, 0.0076 mmol, 1.5 eq.) in DMSO. The reaction mixture was continuously stirred at room temperature. Progress of the reaction was monitored by LC-MS. After complete conversion of folate-nigericin-DBCO, the crude reaction mixture was purified by RP-HPLC, (mobile phase A = 10 mM ammonium acetate, pH = 7; organic phase B = acetonitrile; method: 0% B to 50% B in 35 min at 13 ml/min) and furnished folate-nigericin amide 65% yield. LC-MS (A = 10 mM ammonium bicarbonate, pH = 7; organic phase B = acetonitrile; method: 0% B to 100% B in 12 min) RT = 3.9 min (M + H^+^ = 1728.4).

### Preparation of folate-nigericin-DBCO-miR-34a (Fol-Nig-34a) conjugate

MiRNA duplexes were constructed as previously described^1^. Briefly, a bi-orthogonal click reaction was performed between folate-nigericin-DBCO and azide modified antisense miR-34a (or scramble). Click reaction was performed at a 1:10 molar ratio (azide oligo: folate-nigericin-DBCO) at room temperature in water for eight hours and then cooled to 4°C for four hours. Unconjugated folate-nigericin-DBCO was removed from the reaction using Oligo Clean and Concentrator (Zymo Research) per manufacturer instructions. Conjugation was verified using 15% TAE native polyacrylamide gel electrophoresis (PAGE) and MALDI spectral analysis. After conjugation, the miR-34a sense strand was annealed to the folate-nigericin-miR-34a antisense conjugate. Folate-nigericin-DBCO-miR-34a antisense and miR-34a sense were mixed in an equal molar ratio (1:1, final concentration, 20 uM each) in annealing buffer: 10 mM Tris buffer, pH 7 (Sigma), supplemented with 50 mM NaCl (Sigma), and 1 mM EDTA (Sigma), and incubated at 95°C for five min, cooled slowly to room temperature and then stored at –80°C. This compound is referred to as Fol-Nig-34a. For siRNA studies, the previously reported oligonucleotides targeting Ras homolog gene family, member A (siRhoA)^18^ were used: sense 5’-GACAUGCUUGCUCAUAGUCTT-3’, antisense 3’-TTCUGUACGAACGAGUAUCAG-5’. For cellular uptake experiments, an Atto-647N fluorophore was attached to the 5′ end of the miR-34 antisense oligo and was used in the annealing step generating the following compounds: Fol-34a-Atto647N, and Fol-Nig-34a-Atto647N.

### Stability assay in serum

Stability assays were performed as previously described^1^. Briefly, Fol-Nig-34a conjugates were incubated in 50% fetal bovine serum (Sigma) in water at 37°C for the indicated times. RNA samples were collected and analyzed using 15% TAE PAGE. The reducing agent tris(2-carboxyethyl)phosphine (TCEP) was used as a positive control for nigericin reduction (5mM, 20 min at room temperature).

### Preparation of folate-cyanide 5 (Fol-Cy5) dye conjugate

Folate-cyanide 5 synthesis was performed as previously described^15^. DIPEA (0.0004 g, 0.0032 mmol, 2 eq.) was added dropwise to a stirred solution of folate-EDA (0.0008 g, 0.0016 mmol, 1 eq.) and NHS-Cyanide 5 (Cy5, 0.001 g, 0.001 mmol, 1.1 eq.) in DMSO. The reaction mixture continued to be stirred at room temperature. Progress of the reaction was monitored by LC-MS. After complete conversion of folate-Cy5, the crude reaction mixture was purified by RP-HPLC. (mobile phase A = 10 mM ammonium acetate, pH = 7; organic phase B = acetonitrile; method: 0% B to 50% B in 35 min at 13 ml/min) and furnished folate-Cy5 dye conjugate in 85% yield. LC-MS (A = 10 mM ammonium bicarbonate, pH = 7; organic phase B = acetonitrile; method: 0% B to 50% B in 12 min) RT = 2.40 min (M +H^+^ = 949.2).

### Preparation of folate-nigericin-cyanide 5 (Fol-Nig-Cy5) dye conjugate

Folate-nigericin-cyanide 5 synthesis was performed by adding DIPEA (0.0004 g, 0.0032 mmol, 2. eq.) dropwise to a stirred solution of Folate-nigericin (0.0023 g, 0.0016 mmol, 1 eq.) and NHS-Cy5 (0.001 g, 0.0016 mmol, 1 eq.) in DMSO (Supplemental Figure 1). The reaction mixture continued with stirring at room temperature. Progress of the reaction was monitored by LCMS. After complete conversion of Folate-nigericin- Cy5, the crude reaction mixture was purified by RP-HPLC, (mobile phase A = 10 mM ammonium acetate, pH = 7; organic phase B = acetonitrile; method: 0% B to 50% B in 35 minutes at 13 ml/min) and furnished folate-nigericin-Cy5 65% yield. LC-MS (A = 10 mM ammonium bicarbonate, pH = 7; organic phase B = acetonitrile; method: 0% B to 100% B in 12 minutes) RT = 7.05 min (M + H^+^ = 1906.0).

### Flow Cytometry

Flow cytometry was performed as previously described^1^. FR positive human MB- 231 cells or KB carcinoma cells, or FR negative A549 cells were detached by trypsinization, washed twice in ice-cold phosphate buffered saline (PBS; pH 7.4) and resuspended to a density of 1×10^7^ cells/mL in serum free medium. Cell viability was determined by trypan blue exclusion and cells were only used if the viability of cells was >80%. Next, flow cytometric analyses were performed following standard protocols. One-hundred µL of the cell suspension was incubated with Fol-34a-Atto647N, Fol-Nig-34a-Atto647N, or folate-cyanide 5 (Fol-Cy5) (50nM) in the absence or presence of 10- to 100-fold molar excess of folate glucosamine conjugate. For detection of PDL1, cells were treated with the indicated folate conjugates, or transfected with miR-34a mimics as in^1^. Forty-eight hours later cells were collected and incubated with a 1:100 dilution of anti-PDL1 antibody (Cell Signaling 13684; 0.5 µL/50 µL cold PBS) for 1 h on ice. Unbound antibody was removed by washing cells with ice-cold PBS. Cells were then incubated with 0.5 µL of Alexa Fluro 647 secondary antibody (Thermo Fisher, A-21244) for detection. Cells were incubated at 4°C for 20 min and washed twice with ice cold PBS and analyzed by flow cytometry using LSR Fortessa flow cytometer (BD Biosciences, San Jose, CA, USA). Data was analyzed using FlowJo software v10 (Tree Star, Inc, Ashland, Ore).

### Live Cell Imaging

Briefly, eight-well chambered slides with a glass cover slip bottom (Lab-TekTM Chambered Coverglass, Thermo Fisher Scientific, Denmark) were pre-treated with Poly-D-Lysine (0.1 mg/mL ; Sigma-Aldrich) for five min, washed with PBS and air-dried for five min. KB or MB-231 cells were plated onto prepared slides one day before the experiment at 2×10^4^ cells/well and maintained in RPMI 1640 medium, no folic acid (Life Technologies) supplemented with 10% fetal bovine serum (Sigma), 100 U/mL penicillin and 100 ug/mL streptomycin (Hyclone, GE Healthcare Life Sciences) at 37°C in 5% CO_2_. The slides were placed in a Nikon A1Rsi confocal microscope with a resonant scanner and piezo z-drive (Nikon Instruments Inc.), 20X objective Plan Apo 0.75 NA, equipped with a Tokai hit live imaging chamber (INU-TIZ-F1; Tokai Hit Corp., Japan) with temperature set to 37°C and continuous bubbling of 5% CO_2_ into water. Image acquisition using NIS-elements software 4.5 (Nikon Instruments Inc., Japan) on a single focal plane was conducted for 3 h after addition of Fol-Cy5, Fol-Nig-Cy5, Fol-34a-Atto647N or Fol-Nig-34a-Atto647N conjugates (50 nM). Images were further analyzed using ImageJ 2.0.0 (NIH).

### Segmentation and quantification of subcellular vesicles

Segmentation and quantification of subcellular vesicles was performed using the segmentation and quantification of subcellular shapes (Squassh) protocol^19,20^ in the Mosaic plugin in ImageJ V2.0 (NIH) using the following segmentation parameters: regularization: 0.1, minimum object intensity: 0.160.

### Live cell proliferation assay

KB cells were transduced with 1% V/V NucLight Red Lentivirus (Essen Bioscience, Cat. No 4476), seeded on 96-well plates (Corning 3603; 2,000 cells/well) and incubated at 37°C with 5% CO_2_ overnight. The next day, new medium containing different concentrations of the following reagents was added to the wells: vehicle (DMSO), nigericin, and folate nigericin. The plate was placed in an Icucyte S3 instrument (Essen Bioscience), each well was imaged using standard mode scan with a 10X objective, phase and red filter acquisition, and imaged every four hours for 24 h. Each treatment consisted of three wells with five fields of view per well. Cell proliferation was quantified by counting the number of fluorescent nuclei over 24 h using the Incucyte S3 Software (Essen Bioscience). Data was exported and analyzed using Graphpad Prism v.7. Cell growth is represented as confluence (%) relative to time 0. To determine the dose that causes growth inhibition of at least 50% (GI_50_), the 24-hour data points were selected and normalized to vehicle treated cells.

### Argonaute2 (Ago2) knockdown

GeneSolution siRNAs (Qiagen; 1027416) were used to knockdown Ago2 following the manufacturer’s instructions. Briefly, MB-231 cells were transfected with 25 pmol siRNAs using 7.5 uL of Lipofectaime RNAimax (Life Technologies) in 6 wells (80% confluency). Protein knockdown was measured after 48 hours. siLuc2 was used as a negative control.

### RNA immunoprecipitation assay

RNA precipitation assays were performed as previously described^21,22^. Briefly, MB-231 cells transfected with a miR-34a mimic were used as control. MB-231 cells (7×10^5^ cells/well) were seeded into 6-well plates and maintained overnight in RPMI 1640 medium, no folic acid (Life Technologies) supplemented with 10% fetal bovine serum (Sigma), 100U/mL penicillin and 100ug/mL streptomycin (Hyclone, GE Healthcare Life Sciences) at 37°C in 5% CO_2_. The next day, medium was replaced with 800 μL of fresh complete medium. Transfection of miR-34a mimic (Ambion, final concentration 6nM) was performed with Lipofectamine RNAimax (Life Technologies) by mixing 100 μL serum free medium (SFM) containing mir-34a mimic (Ambion) and 100 μL SFM with 7.5 μL of Lipofectamine RNAimax (Life Technologies), incubated at room temperature for 5 min and then added to the wells. Cells were incubated for 4 h at 37°C in 5% CO_2_ and media was replaced with 2 mL of fresh complete medium. For FolamiR-34a or Fol-Nig-34a treatments, MB-231 cells (8×10^5^ cells/well) in complete medium were seeded into 6-well plates containing FolamiR-34a or Fol-Nig-34a diluted in 1 mL serum free RPMI medium to a final concentration of 100 nM. After 24 h cells were put on ice, washed twice with ice cold PBS and immersed in 1 mL PBS. Cells were cross-linked at 400 mJ per cm^2^ and then again at 200 mJ per cm^2^ in an UV cross-linker (XL-1000, SpectroLinker). Cells were scraped directly into lysis buffer (1× PBS, 1% vol/vol NP40 substitute, 0.5% wt/vol sodium deoxycholate and 0.1% wt/vol SDS). Next, the lysates were centrifuged in a prechilled tabletop ultracentrifuge at 21,000 g for 20 min at 4°C. Cleared lysates were incubated with Dynabead Protein A beads (Life Technologies) prepared with 2A8 anti-AGO^21,23^ (Millipore, MABE56) or normal mouse IgG (Millipore, 12–371) for 90 min at 4°C, washed and resuspended as previously described^21^. RNA was isolated using Qiazol reagent (Qiagen) followed by ethanol precipitation. Reverse transcription was performed using the miRscript II RT kit (Qiagen) using the HiSpec buffer. Real-time PCR was conducted using miScript SYBR Green PCR Kit (Qiagen) with the following primers: RNU6B (non-target RNA, Qiagen MiScript primer assay), *let-7b-1* (positive control, Qiagen MiScript primer assay) and mir-34a (Qiagen MiScript primer assay). Real time PCR data was analyzed using the 2^−▵▵Ct^ method^24,25^ and expressed as fold enrichment.

### Protein isolation and western blotting

MB-231 cells were collected in RNA Later (Life Technologies) and stored at –80°C. Frozen cells were placed in 2 mL collection tubes containing radioimmunoprecipitation assay buffer (1 X phosphate-buffered saline, 1% NP-40, 0.5% sodium deoxycholate, 0.1% sodium dodecyl sulfate, 1 X protease inhibitor). Cells were incubated for 10 min on ice and centrifuged for 10 min at 4°C (20,000 g), supernatant was collected, 50 ug of protein was resolved on 12% TGX gels (Bio-Rad) and transferred to polyvinylidene difluoride (PVDF) membranes. Revert Total Protein Staining kit (Licor) was used as protein loading control following the manufacturer’s instructions. 2A8 anti-AGO (1:1000, Millipore, MABE56) primary antibody was used for Argonaute 2 detection. Protein detection was performed following incubation with secondary antibodies (Licor), membranes were washed and signal was acquired using Licor Odyssey CLX (Licor).

## Acknowledgements

We thank members of the Kasinki lab for critical review of the manuscript and both the Kasinski and Low Laboratories for insightful discussion. This work was supported by NCI R01CA226259 (ALK) and NCI R01CA205420 (ALK). EAO was supported by a Bilsland Dissertation Fellowship from Purdue University. AMA was supported by an Andrew’s Fellowship from Purdue University. The authors gratefully acknowledge support from the Purdue University Center for Cancer Research, NIH grant P30 CA023168, and the Purdue University Center for Cancer Research Small Grants Program.

## References

1. Orellana, EA, Tenneti, S, Rangasamy, L, Lyle, LT, Low, PS and Kasinski, AL (2017). FolamiRs: Ligand-targeted, vehicle-free delivery of microRNAs for the treatment of cancer. Science Translational Medicine 9: eaam9327.

2. Zuckerman, JE, Gritli, I, Tolcher, A, Heidel, JD, Lim, D, Morgan, R, et al. (2014). Correlating animal and human phase Ia/Ib clinical data with CALAA-01, a targeted, polymer-based nanoparticle containing siRNA. Proceedings of the National Academy of Sciences 111: 11449–11454.

3. Wang, H, Jiang, Y, Peng, H, Chen, Y, Zhu, P and Huang, Y (2015). Recent progress in microRNA delivery for cancer therapy by non-viral synthetic vectors. Adv. Drug Deliv. Rev. 81: 142–160.

4. Orellana, EA and Kasinski, AL (2017). No vehicle, no problem. Oncotarget 8: 96470–96471.

5. Shete, HK, Prabhu, RH and Patravale, VB (2014). Endosomal Escape: A Bottleneck in Intracellular Delivery. j. nanosci. nanotech. 14: 460–474.

6. Vlashi, E, Kelderhouse, LE, Sturgis, JE and Low, PS (2013). Effect of folate-targeted nanoparticle size on their rates of penetration into solid tumors. ACS Nano 7: 8573–8582.

7. Yang, J, Chen, H, Vlahov, IR, Cheng, J-X and Low, PS (2006). Evaluation of disulfide reduction during receptor-mediated endocytosis by using FRET imaging. Proc. Natl. Acad. Sci. U.S.A. 103: 13872–13877.

8. Yang, J, Chen, H, Vlahov, IR, Cheng, J-X and Low, PS (2007). Characterization of the pH of folate receptor-containing endosomes and the rate of hydrolysis of internalized acid-labile folate-drug conjugates. J. Pharmacol. Exp. Ther. 321: 462–468.

9. Varkouhi, AK, Scholte, M, Storm, G and Haisma, HJ (2011). Endosomal escape pathways for delivery of biologicals. Journal of Controlled Release 151: 220–228.

10. Lundberg, P, El-Andaloussi, S, Sütlü, T, Johansson, H and Langel, Ü (2007). Delivery of short interfering RNA using endosomolytic cell-penetrating peptides. FASEB Journal 21: 2664–2671.

11. Oliveira, S, van Rooy, I, Kranenburg, O, Storm, G and Schiffelers, RM (2007). Fusogenic peptides enhance endosomal escape improving siRNA-induced silencing of oncogenes. International Journal of Pharmaceutics 331: 211–214.

12. Erazo-Oliveras, A, Muthukrishnan, N, Baker, R, Wang, T-Y and Pellois, J-P (2012). Improving the Endosomal Escape of Cell-Penetrating Peptides and Their Cargos: Strategies and Challenges. Pharmaceuticals 2012, Vol. 5, Pages 1177–1209 5: 1177–1209.

13. Behr, J-P (1997). The proton sponge: A trick to enter cells the viruses did not exploit 51, 3pp: pp 34–36.

14. Oliveira, S, Fretz, MM, Høgset, A, Storm, G and Schiffelers, RM (2007). Photochemical internalization enhances silencing of epidermal growth factor receptor through improved endosomal escape of siRNA. Biochimica et Biophysica Acta (BBA) – Biomembranes 1768: 1211–1217.

15. Rangasamy, L, Chelvam, V, Kanduluru, AK, Srinivasarao, M, Bandara, NA, You, F, et al. (2018). New Mechanism for Release of Endosomal Contents: Osmotic Lysis via Nigericin-Mediated K^+^/H^+^Exchange. Bioconjugate Chem. 29: 1047–1059.

16. Steinrauf, LK, Pinkerton, M and Chamberlin, JW (1968). The structure of nigericin. Biochem. Biophys. Res. Commun. 33: 29–31.

17. Graven, SN, Estrada-O, S and Lardy, HA (1966). Alkali metal cation release and respiratory inhibition induced by nigericin in rat liver mitochondria. Proc. Natl. Acad. Sci. U.S.A. 56: 654–658.

18. Sun, H-W, Tong, S-L, He, J, Wang, Q, Zou, L, Ma, S-J, et al. (2007). RhoA and RhoC –siRNA inhibit the proliferation and invasiveness activity of human gastric carcinoma by Rho/PI3K/Akt pathway. World Journal of Gastroenterology : WJG 13: 3517–3522.

19. Paul, G, Cardinale, J and Sbalzarini, IF (2013). Coupling image restoration and segmentation: A generalized linear model/bregman perspective. International Journal of Computer Vision 104: 69–93.

20. Rizk, A, Paul, G, Pietro Incardona, Bugarski, M, Mansouri, M, Niemann, A, et al. (2014). Segmentation and quantification of subcellular structures in fluorescence microscopy images using Squassh. Nature Protocols 9: 586–596.

21. Moore, MJ, Zhang, C, Gantman, EC, Mele, A, Darnell, JC and Darnell, RB (2014). Mapping Argonaute and conventional RNA-binding protein interactions with RNA at single-nucleotide resolution using HITS-CLIP and CIMS analysis. Nature Protocols 9: 263.

22. Moore, MJ, Scheel, TKH, Luna, JM, Park, CY, Fak, JJ, Nishiuchi, E, et al. (2015). miRNA–target chimeras reveal miRNA 3′-end pairing as a major determinant of Argonaute target specificity. Nat Commun 6: 8864.

23. Nelson, PT, de Planell-Saguer, M, Lamprinaki, S, Kiriakidou, M, Zhang, P, O’Doherty, U, et al. (2007). A novel monoclonal antibody against human Argonaute proteins reveals unexpected characteristics of miRNAs in human blood cells. RNA 13: 1787–1792.

24. Livak, KJ and Schmittgen, TD (2001). Analysis of Relative Gene Expression Data Using Real-Time Quantitative PCR and the 2− ΔΔCT Method. Methods 25: 402–408.

25. Yuan, JS, Reed, A, Chen, F and Stewart, CN (2006). Statistical analysis of real-time PCR data. BMC Bioinformatics 7: 85.

26. Marks, IS, Kang, JS, Jones, BT, Landmark, KJ, Cleland, AJ and Taton, TA (2011). Strain-promoted ‘click’ chemistry for terminal labeling of DNA. Bioconjugate Chem. 22: 1259–1263.

27. Parker, N, Turk, MJ, Westrick, E, Lewis, JD, Low, PS and Leamon, CP (2005). Folate receptor expression in carcinomas and normal tissues determined by a quantitative radioligand binding assay. Analytical Biochemistry 338: 284–293.

28. Liu, J, Valencia-Sanchez, MA, Hannon, GJ and Parker, R (2005). MicroRNA-dependent localization of targeted mRNAs to mammalian P-bodies. Nat. Cell Biol. 7: 719.

29. Leung, AKL, Calabrese, JM and Sharp, PA (2006). Quantitative analysis of Argonaute protein reveals microRNA-dependent localization to stress granules. Proc. Natl. Acad. Sci. U.S.A. 103: 18125–18130.

30. Nair, JK, Willoughby, JLS, Chan, A, Charisse, K, Alam, MR, Wang, Q, et al. (2014). Multivalent N-acetylgalactosamine-conjugated siRNA localizes in hepatocytes and elicits robust RNAi-mediated gene silencing. J. Am. Chem. Soc. 136: 16958–16961.

31. Matsuda, S, Keiser, K, Nair, JK, Charisse, K, Manoharan, RM, Kretschmer, P, et al. (2015). siRNA conjugates carrying sequentially assembled trivalent Nacetylgalactosamine linked through nucleosides elicit robust gene silencing in vivo in hepatocytes. ACS chemical biology 10: 1181–1187.

32. Rajeev, KG, Nair, JK, Jayaraman, M, Charisse, K, Taneja, N, O’Shea, J, et al. (2015). Hepatocyte-specific delivery of siRNAs conjugated to novel non-nucleosidic trivalent Nacetylgalactosamine elicits robust gene silencing in vivo. Chembiochem 16: 903–908.

33. Sehgal, A, Barros, S, Ivanciu, L, Cooley, B, Qin, J, Racie, T, et al. (2015). An RNAi therapeutic targeting antithrombin to rebalance the coagulation system and promote hemostasis in hemophilia. Nat. Med. 21: 492–497.

34. Fitzgerald, K, White, S, Borodovsky, A, Bettencourt, BR, Strahs, A, Clausen, V, et al. (2017). A Highly Durable RNAi Therapeutic Inhibitor of PCSK9. N. Engl. J. Med. 376: 41–51.

